# Visual search training benefits from the integrative effect of enhanced covert attention and optimized overt eye movements

**DOI:** 10.1101/2022.01.03.474790

**Authors:** Qi Zhang, Zhibang Huang, Liang Li, Sheng Li

**Affiliations:** School of Educational Science, Minnan Normal University; School of Psychological and Cognitive Sciences, Peking University; Beijing Key Laboratory of Behavior and Mental Health, Peking University; PKU-IDG/McGovern Institute for Brain Research, Peking University; Key Laboratory of Machine Perception (Ministry of Education), Peking University; The Brain Science Center, Beijing Institute of Basic Medical Sciences

**Keywords:** conjunction search, saccade, search initiation time, N2pc.

## Abstract

Training serves as an effective approach to improve visual search performance when the target does not automatically pop out from the distractors. In the present study, we trained participants on a conjunction visual search task and examined the training effects in behavior and eye movement. The results of Experiments 1 to 4 showed that training improved behavioral performance and reduced the number of saccades and overall scanning time. Training also increased the search initiation time before the first saccade and the proportion of trials in which the participants correctly identified the target without any saccade, but these effects were modulated by stimulus’ parameters. In Experiment 5, we simultaneously recorded eye movements and EEG signals and the results revealed significant N2pc components after the stimulus onset (i.e., stimulus-locked) and before the first saccade (i.e., saccade-locked) when the search target was the trained one. These N2pc components can be considered as the neural signatures for the enhanced covert attention to the trained target. Together with the training-induced increase in functional visual field, these mechanisms could support the beneficial effects of increased search initiation time and reduced number of saccades. These findings suggest that visual search training enhanced covert attention to target and optimized overt eye movements to facilitate search performance.

---

Effective identification of a search target in a complex visual environment relies on multiple factors. To reveal the complex mechanisms underlying this simple behavior, investigators have developed a variety of visual search tasks in which participants look for a pre-defined target among various distractors (Duncan & Humphrey, 1989; Theeuwes, 1992; Treisman & Gelade, 1980). According to the patterns of searching behavior, a search task is called parallel if a single feature that could easily pop-out from homogeneous distractors defines the target. As a result, reaction time (RT) in parallel search is independent of the number of distractors (Bergen & Julesz, 1983; Duncan, 1989; Egeth et al., 1972). In contrast, in a conjunction visual search task in which the target is defined by combination of two features and may share one of the two features with distractors, the task is serial as it requires shifts of attentional focus among search items (Treisman & Gelade, 1980). Consequently, reaction time in serial visual search increases as the number of distractors increase (Treisman & Paterson, 1984).

As we live in a diverse and dynamic world, experience-dependent adjustments of visual functions are necessary. Training on visual search tasks in laboratory is a useful approach to examine the experience-dependent visual behavior. According to the influential feature integration theory (Treisman & Gelade, 1980), a unitary detector of a conjunction of features could emerge after extended practice on the stimuli with that conjunction. The emergence of the unitary detector would lead to a change of search mode from serial to parallel. However, the unitary detector also implies that the training effect would not be transferred to the stimuli in which one of the conjunctive features is changed. Another important theory of attention, the guided search model, proposed that visual search process is simultaneously serial and parallel, and selection history could serve as a source of information to guide search process (Wolfe, 1994, 2021). The guided search model also suggests that, when the same target is searched consecutively for many times, higher attention priority would be assigned to it in the subsequent search. Particularly, the participants could learn to pay attention to the specific features of a conjunction target more effectively, making the transfer of the training effect possible if a stimulus retains only a fraction of the conjunctive features (Andersen et al., 2008).

In recent years, various forms of visual search paradigms had been adopted in visual search training, including visual search of shape (Qu et al., 2017; Sigman & Gilbert, 2000), orientation (An et al., 2012), and letter (Bueichekú et al., 2016, 2019), as well as various conjunction visual search tasks (Su et al., 2014; Frank et al., 2016; Reavis et al., 2016, 2018). These studies all demonstrated that training can substantially improve visual search performance. Among them, Su et al. (2014) demonstrated a feature-based attention enhancement mechanism rather than a unitization mechanism. Transfer of the training effects to untrained stimuli was also found in different tasks (Sirenteanu & Rettenbach, 2000; Sireteanu & Rettenbach, 1995). These results agreed with the hypothesis of the guided search model (Wolfe, 1994, 2021). To reveal the neural mechanisms behind the observed improvement, neuroimaging techniques had been adopted in other investigations. For example, the increased early sensory-evoked N1 component was found after visual search training (Clark et al., 2015). Particularly, Qu et al. (2017) presented strong evidence that after training, trained target stimulus that was physically non-salient could capture attention as indicated by the N2 posterior contralateral (N2pc) ERP component. Studies that used conjunctive stimuli defined by the spatial configuration of two elements also revealed training-induced performance improvement (Reavis et al., 2016, 2018).

Using fMRI, these studies found that conjunction learning in visual search led to an increase in target-evoked activity relative to distractor-evoked activity even when participants passively viewed the trained stimuli while performing an unrelated, attention-demanding task (Reavis et al., 2016). Overall, the literature suggested that training on visual search could facilitate the detection of target on the level of object or feature.

The guided search model proposed that the search process is simultaneously serial and parallel (Wolfe, 2021). Correspondingly, visual search is a process that is generally recruits both covert attention and overt attention (Hoffman & Subramaniam, 1995; Li et al., 2021; Moore & Fallah, 2001; Posner, 1980). However, while the parallel search is generally associated with covert attention, the serial search could involve both modes. As suggested by the previous literature, training could facilitate the search performance by the transformation from serial search to parallel search. Indeed, attentional capture by previously trained target stimulus could be a hint for this transformation process (Qu et al., 2017). However, how training concurrently modulates the covert and overt aspects of attention in visual search remained a less investigated topic. The above-mentioned studies had adopted paradigms in which participants were required to maintain fixation during the search task and therefore focused more on the covert aspect of attentional processing. In natural viewing conditions, the optimal efficiency for identifying a search target would be achieved by the combination of enhanced covert attention and improved overt eye movements. Mandatory fixation during the search process may not be sufficient to reveal the integrative effect of training in both of them. In the present study, we used a free viewing paradigm and recorded participants’ eye movements with an eye tracker to avoid this problem.

In the free viewing paradigm in which the participants can shift their gazes among search items, the duration of eye movement can be decomposed into three epochs (Malcolm & Henderson, 2009): search initiation time as the fixational period before the first saccade, scanning time as the period from the first saccade until the beginning of the last fixation, and verification time as the duration of the last fixation until the participants’ response. Given our hypothesis that training would induce an integrative effect of covert and overt attention, we took the advantage of this approach and examined whether the specific epochs and the related measurements of covert and overt attention were modulated by visual search training. Specifically, we examined the proportion of correct trials in which no saccade (zero- saccade trials) or only one on-target saccade (single on-target-saccade trials) was made before response. These two measurements were tightly associated with the covert attention that may contribute to the training-induced transformation from serial to parallel search. In addition, we attempted to measure the functional visual field (FVF) in these two types of trials as an indicator of the spatial scope of attention that we believed training effect might have taken place (Wolfe, 2021; Wu & Wolfe, 2022). Stimulus parameters (e.g., stimulus size and set size) and crowding effect that are related to attentional processing were also under our consideration.

Further, by simultaneously recording eye movements and EEG signals, we could also reveal the neural signatures of covert attention that were accompanied with the training effects in overt eye movements. Particularly, we examined saccade-locked N2pc component (Huber-Huber et al., 2016; Talcott & Gaspelin, 2021; Weaver et al., 2017) along with the classical stimulus-locked N2pc that measured the stimulus-induced attentional bias (Eimer, 2014; Luck & Hillyard, 1994; Hickey et al., 2006). New insights into the training-induced integrative effect of covert attention and overt eye movements could be gained given this combinatorial approach.

## Overview of the Study

We conducted four behavioral experiments (Experiments 1 to 4) and an EEG experiment (Experiment 5). Eye movement data were recorded in all experiments. Search arrays in the experiments were composed of line segments with two defining features (i.e., color and orientation, Figure 1A-C). In Experiments 1 to 3, we trained participants in a conjunction visual search task and tested their performance before and after the training. Experiment 1 was conducted to establish a standard paradigm for evaluating the training effects. In this experiment, we examined the transfer of the training effect to stimuli that shared one of the two features with the trained target. By changing the set size and stimulus size in Experiments 2 and 3, we also investigated the roles of stimulus parameters on the training effect. Particularly, we examined the influence of crowding when the set size was increased. In Experiment 4, the training was replaced with a two-hour rest period to establish a baseline performance without training that would be attributed to the practice effect of the training protocol. The results showed the behavioral improvement was accompanied with reduced fixation number and hence scanning time (Experiments 1 to 3), as well as increased proportion of zero-saccade correct trials and search initiation time (Experiment 1). The practice effect was not sufficient to generate the training-induced behavioral improvement (Experiment 4). In Experiment 5, we replicated these behavioral and eye movement findings when EEG signals were also simultaneously recorded. The results revealed enlarged spatial scope of attention as indexed by the functional visual field and identified stimulus-locked and saccade-locked N2pc components as the neural signatures for the training-induced enhancement in covert attention to the trained target.

**Figure 1.**
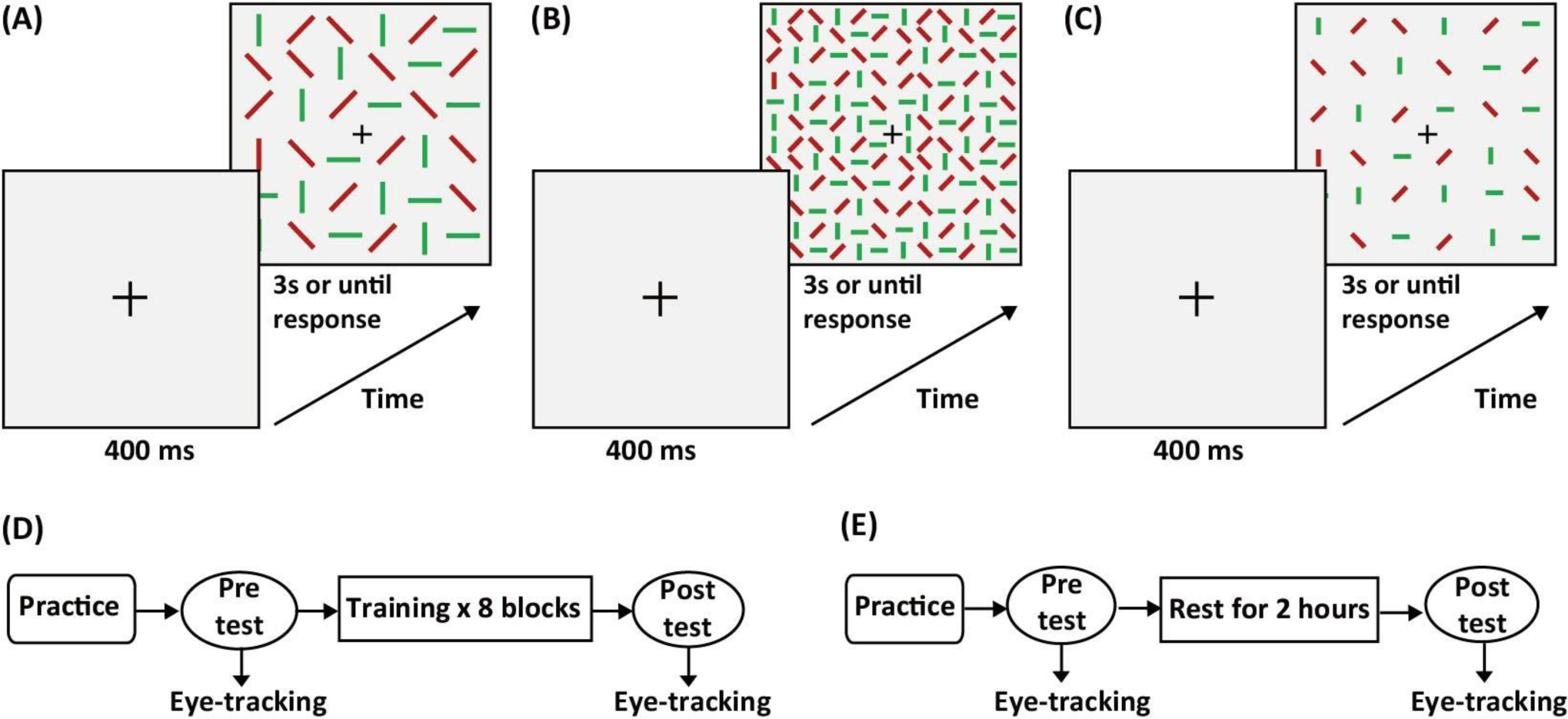
Stimuli and typical trials of the conjunction visual search task in (A) Experiment 1, (B) Experiments 2 and 4, and (C) Experiment 3. (D) Procedure in Experiments 1, 2, and 3. (E) Procedure in Experiment 4.

We totally recruited 80 participants with 15 participants for Experiments 1 to 4 and 20 participants for Experiment 5. We conducted a priori power analyses with G*Power 3.0 (Faul et al., 2007). The analyses suggested that 15 participants are required for each experiment to detect a behavioral training effect on visual search (*F* tests: repeated-measures ANOVA, within factor, expected effect size *η^2^* = 0.298, power = 0.9). The expected effect size was chosen based on previous studies that adopted similar conjunction visual search training paradigm (Su et al., 2014, Experiment 3a). We recruited 20 participants in Experiment 5 to increase the power for detecting ERP effects, because EEG signals have lower signal-to-noise ratio and 15 participants were estimated based on behavioral results. The experiments were approved by the local ethics committee. The participants provided written informed consent prior to the experiment in accordance with the Declaration of Helsinki and were paid for their participation.

## Experiment 1

In Experiment 1, we trained participants on a conjunction visual search task. We examined the training effects on the trained target and two stimuli that either shared color or orientation feature with the trained target when they served as target in the pretest and posttest sessions. We also calculated transfer indices to quantify the transfer of training effects to the stimulus that shared color or orientation with the trained target.

### Method

#### Participants

Fifteen right-handed naïve participants (eight females, age range = 18–27 years, mean age = 22.5 years) with normal or corrected-to-normal vision were recruited for the experiment.

#### Apparatus

All stimuli were displayed on a 32-inch Display++ LED monitor (Cambridge Research Systems Ltd, Rochester, Kent, UK) with a refresh rate of 120 Hz and spatial resolution of 1920 × 1080 pixels. The stimuli were presented using Psychtoolbox 3.0 (Brainard, 1997; Pelli, 1997) in MATLAB programming environment (MathWorks, Natick, MA, USA). The participants were positioned 72 cm from the monitor in a dimly lit room. We used a chin rest to fix the participants’ head position. Eye movements were recorded using an EyeLink 1000 plus (SR Resear ch, Ontario, Canada) eye tracker with a sampling rate of 1000 Hz. Gaze position was established using nine-point calibration and validation procedure. Drift correction was performed before each block.

#### Stimuli

Each search array (16.37° × 16.37°) consisted of a central fixation cross and 36 items (1.37° × 0.55°) in a 6 × 6 array (Figure 1A). The items were line segments on a gray background with distinctive colors (red or green) and orientations (0°: horizontal, 90°: vertical, 45°: tilted right from vertical, or 135°: tilted left from vertical). In half of the trials, there was a unique target (i.e., red 90°, denoted as r_90) in the search array and the rest of the items were distractors (randomly chosen from g_0, g_90, r_45, and r_135). In the other half of the trials, there was no target and all items in the search array were distractors.

#### Procedure

The procedure of the experiment is shown in Figure 1D. Participants completed a practice block (100 trials) to familiarize with the task before the pretest session. In the practice block, search arrays consisted of black and white line segments with a black 90° line segment serving as target. In the pretest, participants were tested for three different targets (r_90, r_0, and g_90 with equal number of trials) separately in three blocks with 100 trials in each block. After the pretest, participants conducted eight training blocks with 400 trials in each block. The targets were always r_90 in the training blocks and appeared in a half of the trials. There was a three minutes break after every block and a five minutes break after every two blocks. During the practice and training, participants were offered an optional break every 20 trials to reduce fatigue and boredom. After training, participants completed a posttest that was identical to the pretest. The orders of the blocks in the pretest and posttest sessions were randomized.

A typical trial of the conjunction visual search task is shown in Figure 1A. Each trial began with the presentation of a central fixation cross for 400 ms, followed by a search array that appeared until the key response or the elapse of 3000 ms since its onset. Participants were instructed to fixate on the central fixation before the presentation of the search array and return to the fixation cross as soon as possible after they found the target. They could make eye movements during search period. Participants were required to indicate the presence or absence of the target as accurately and quickly as possible by pressing one of two keys with one of two fingers of the right hand.

#### Data analysis

##### Behavior

Trials with incorrect responses (4.01%), RTs that were faster than 200 ms (0.09%), and latency of first saccade smaller than 80 ms (4.27%), were excluded from analyses. The trials in which the participants failed to respond within 3000 ms were considered as incorrect trials. Both target-present and target-absent trials were included in the analysis. In the calculation of d’, when participants scored perfect performance (i.e., hit rate = 1 or false alarm rate = 0), hit rate was assigned a value of 1- 1/ (2 × N) or false alarm rate was assigned a value of 1/ (2 × N) (N was the total number of target-present or target-absent trials). For the presentation purpose, we refer the trials with the target of r_90, r_0, and g_90 as C+O+, C+O–, and C–O+, respectively (C for color feature, O for orientation feature, + and – indicated that the feature was the same as and different from the trained target, respectively). Repeated-measures two-way analyses of variance (ANOVAs) with session (pretest and posttest) and target type (C+O+, C+O–, and C–O+) as factors were conducted with SPSS (version 20, IBM), and if necessary, the measures were corrected for violation of sphericity assumption by using Greenhouse–Geisser correction. Corrections made for multiple comparisons were also reported.

We defined transfer index (TI) from C+O+ to either C+O– or C–O+ as the mean percent improvement (MPI) of the untrained target (C+O–, C–O+) divided by the MPI of the trained target (C+O+). Therefore, transfer index of 1 would suggest a full transfer of training effect from the trained target to the untrained target. In contrary, transfer index of 0 would imply that there was no transfer of training effect at all. The MPI of each participant and each condition was calculated as (posttest– pretest)/pretest.

##### Eye movement

We divided the correct trials into three categories: zero-saccade trials (no saccade before response), single on-target-saccade trials (only one saccade before response and this saccade was on the target), and other trials. For these other trials with overt eye movement, we defined three epochs for each trial (Malcolm & Henderson, 2009): search initiation time, scanning time, and verification time. Search initiation time was defined as the period from the onset of the search display until the first saccade away from the central fixation. We assumed that search initiation time reflected the time needed to select the first item in the search display for examination. Scanning time was defined as the period from the first saccade until the onset of the last fixation. This epoch reflected the actual search process. We also calculated the number of fixations and mean fixation duration during the scanning period. Verification time was defined as the duration of the last fixation until the participants’ response for those target-present trials in which the last fixation was on the target. This epoch reflected the time to determine whether the fixated item was the target. Segmenting total trial duration into three epochs helped us to elucidate the training effect on these specific search processes.

Two issues need to be clarified for the definition of eye movements. First, a fixation was defined as on target or central cross if its distance from the center of the target or central cross was shorter than 1.5°. Second, we observed that the participants occasionally returned their gaze back to the central fixation to prepare for next trial but this was done before their response to the current trial. Therefore, if there were more than one fixation in a trial, and the last fixation was on the central cross and its start time was within 300 ms from the key response, we removed this last fixation from the analysis and subtracted one from the number of fixations.

Both target-present and target-absent trials were included in the eye movement analysis, with the exception for the proportions the single on-target-saccade trials and the verification time of other trials that were based on the target-present trials. The statistical analyses (i.e., ANOVAs) for the measurements of eye movement were conducted in the same way as for the behavioral measurements.

### Results

#### d’ and RTs

As shown in Figure 2A, the behavioral results of d’ and RTs suggest that training improved visual search performance.

**Figure 2.**
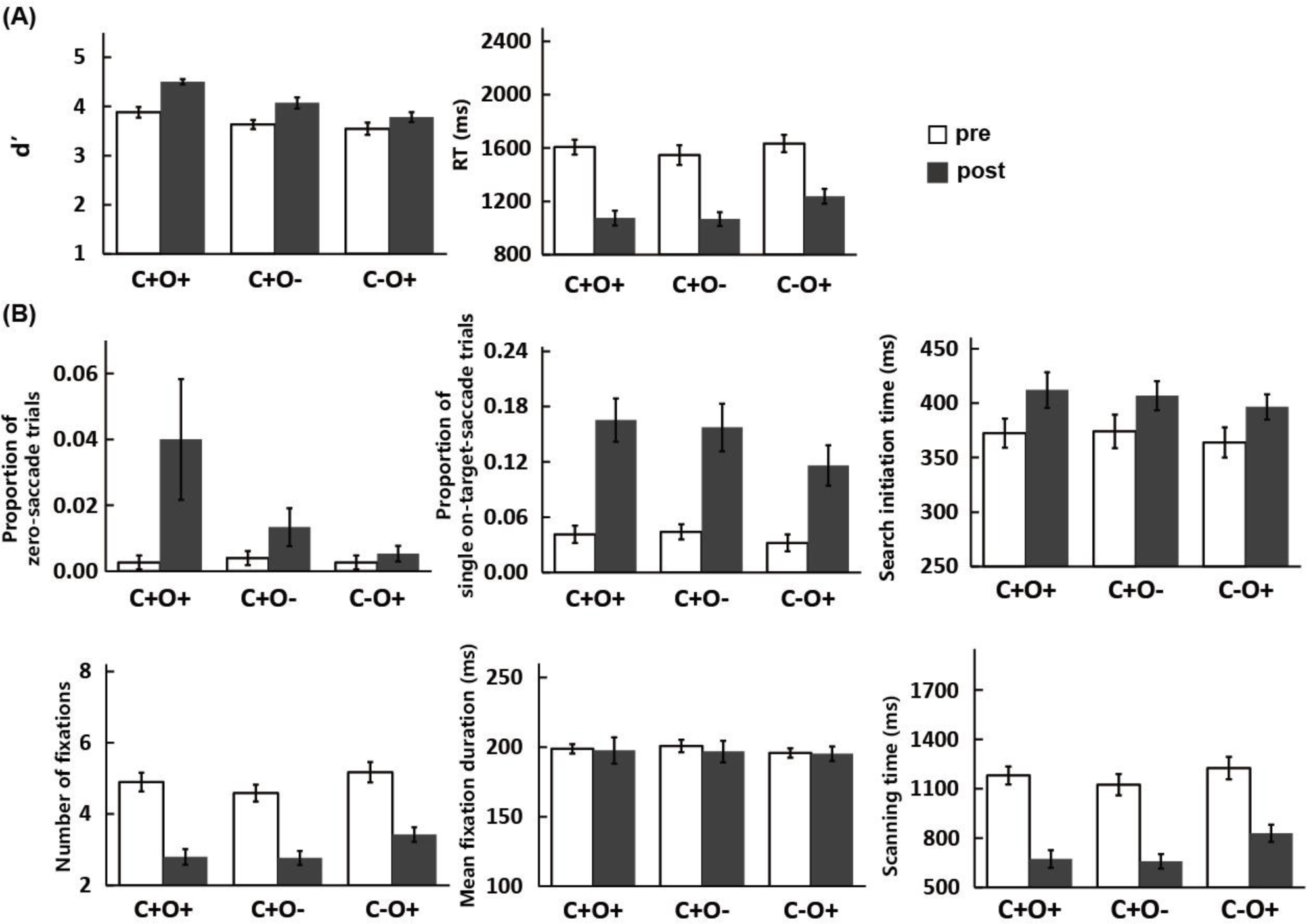
Results of Experiment 1. (A) Behavioral results for d’ and RTs. (B) Eye movement results for the proportion of zero-saccade trials, proportion of single on-target-saccade trials, and other trials’ search initiation time, number of fixations, mean fixation duration, and scanning time. Error bars represent standard errors of the mean across participants.

For d’, we observed significant effects on session (*F*(1,14) = 28.54, *p* < 0.001, *η^2^* = 0.67) and target type (*F*(2,28) = 37.07, *p* < 0.001, *η^2^* = 0.73), but not on their interaction (*F*(2,28) = 1.70, *p* = 0.20, *η^2^* = 0.11). Bonferroni-corrected post hoc tests showed that there were significant differences between C+O+ and C+O– conditions (*p* < 0.001) and between C+O+ and C–O+ conditions (*p* < 0.001). No significant difference was observed between C+O– and C–O+ conditions (*p* = 0.06).

For RTs, we also observed significant effects on session (*F*(1,14) = 82.29, *p* < 0.001, *η^2^* = 0.85) and target type (*F*(2,28) = 4.15, *p* < 0.05, *η^2^* = 0.23), but not on their interaction (*F*(2,28) = 2.77, *p* = 0.08, *η^2^* = 0.16). Bonferroni-corrected post hoc tests showed that there was significant difference between C+O+ and C–O+ condition (*p* < 0.05). No significant difference was observed between C+O+ and C+O– conditions (*p* = 1.00) or between C+O– and C–O+ conditions (*p* = 0.08).

To examine the transfer of learning effect to the stimuli that differed from the trained target in one of the two features, we calculated the transfer indices for C+O–and C–O+ conditions (Table 1). Paired t-tests revealed no significant difference in transfer index between C+O– and C–O+ conditions in d’ (*t*(14) = 1.81, *p* = 0.09, Cohen’s *d* = 0.47) and RTs (*t*(14) = 1.81, *p* = 0.09, Cohen’s *d* = 0.47).

**Table 1.** Transfer indices (mean±SD) from Experiment 1 to Experiment 3.

| **Experiment 1** | **C+O–** | **C–O+** |
| --- | --- | --- |
| d' | 0.91±1.34 | 0.02±1.88 |
| RTs | 0.95±0.39 | 0.74±0.35 |
| Search initiation time | 2.90 ± 6.86 | 1.06 ± 2.10 |
| Number of fixations | 0.97 ± 0.41 | 0.81 ± 0.33 |
| Scanning time | 0.97 ± 0.34 | 0.75 ± 0.31 |
| **Experiment 2** | **C+O–** | **C–O+** |
| d' | 0.64±0.72 | 0.32±0.55 |
| RTs | 0.74±0.48 | 0.30±0.37 |
| Search initiation time | 2.29±3.67 | -0.04±5.07 |
| Number of fixations | 0.82±0.47 | 0.39±0.57 |
| Scanning time | 0.73±0.49 | 0.31±0.38 |
| **Experiment 3** | **C+O–** | **C–O+** |
| d' | -0.01±1.54 | -0.25±1.77 |
| RTs | 0.86±0.80 | 0.77±0.33 |
| Search initiation time | 1.51±8.63 | -0.65±4.23 |
| Number of fixations | -2.10±11.14 | -0.39±4.47 |
| Scanning time | 0.05±2.84 | 0.35±1.46 |

#### Eye movement

The eye movement results are shown in Figure 2B. The trials were grouped as zero-saccade trials, single on-target-saccade trials, and other correctly responded trials.

For the proportions of the zero-saccade trials, there was a significant effect on session (*F*(1,14) = 5.42, *p* < 0.05, *η^2^* = 0.28). No significant effect on target type (*F*(1.09,15.20) = 3.11, *p* = 0.10, *η^2^* = 0.18) or their interaction (*F*(1.17,16.32) = 3.46, *p* = 0.08, *η^2^* = 0.20) was observed.

For the proportions the single on-target-saccade trials, there was also a significant effect on session (*F*(1,14) = 45.67, *p* < 0.001, *η^2^* = 0.77). No significant effect on target type (*F*(2,28) = 2.29, *p* = 0.12, *η^2^* = 0.14) or their interaction (*F*(2,28) = 1.23, *p* = 0.31, *η^2^* = 0.08) was observed.

For other correctly responded trials, training significantly increased the search initiation time and reduced the number of fixations and scanning time. For the search initiation time, there was a significant effect on session (*F*(1,14) = 5.74, *p* < 0.05, *η^2^* = 0.29). No significant effect on target type (*F*(2,28) = 1.81, *p* = 0.18, *η^2^* = 0.11) or their interaction (*F*(2,28) = 0.19, *p* = 0.83, *η^2^* = 0.01) was observed. For the number of fixations, there were significant effects on session (*F*(1,14) = 106.63, *p* < 0.001, *η^2^* = 0.88) and target type (*F*(2,28) = 8.18, *p* < 0.01, *η^2^* = 0.37). No significant effect on their interaction was observed (*F*(2,28) = 1.28, *p* = 0.29, *η^2^* = 0.08). Bonferroni-corrected post hoc tests showed that there were significant differences between C+O+ and C–O+ conditions (*p* < 0.05) and between C+O– and C–O+ conditions (*p* < 0.05). No significant difference was observed between C+O+ and C+O– conditions (*p* = 0.93). For scanning time, there were significant effects on session (*F*(1,14) = 96.69, *p* < 0.001, *η^2^* = 0.87) and target type (*F*(2,28) = 5.36, *p* < 0.05, *η^2^* = 0.28). No significant effect on their interaction was observed (*F*(2,28) = 2.20, *p* = 0.13, *η^2^* = 0.14). Bonferroni-corrected post hoc tests showed that there were significant differences between C+O+ and C–O+ conditions (*p* < 0.05) and between C+O– and C–O+ conditions (*p* < 0.05). No significant difference was observed between C+O+ and C+O– conditions (*p* = 1.00). The mean fixation duration and verification time were not significantly modulated by training (*ps* > 0.10).

For the measurements of eye movement that were significantly modulated by training, we calculated their transfer indices (Table 1). Paired t-tests revealed no significant difference in transfer index between C+O– and C–O+ conditions in the search initiation time (*t*(14) = 0.99, *p* = 0.34, Cohen’s *d* = 0.25), number of fixations (*t*(14) = 1.29, *p* = 0.22, Cohen’s *d* = 0.33), and scanning time (*t*(14) = 1.84, *p* = 0.09, Cohen’s *d* = 0.47) of the other correctly responded trials. For the proportions of the zero-saccade trials and the single on-target-saccade trials, the calculation of the transfer index was not possible due to the existence of zero values in the pretest sessions of individual participants.

### Discussion

The behavioral results in Experiment 1 demonstrated that training on the conjunction visual search task significantly improved search performance of the participants as they identified the target faster and more accurate. The eye movement results revealed that one of the sources for decreased RTs after training was the reduction in the number of fixations (or saccades). In contrary, mean fixation duration and verification time remained unchanged after training, indicating that the processing time for each fixated item was not affected by training.

The behavioral results showed that the training effect was not specific to the trained target stimulus, but also transferred to the stimuli that shared one of the two features with the trained target (i.e., C+O– and C–O+), as was evident with the significant session effect and the absence of its interaction with target type. We calculated the transfer indices for the two untrained conditions. The statistics revealed only trends of significance (*ps* = 0.09, TIs were larger for C+O–) between the two conditions for d’, RTs, and scanning time, indicating a potential larger transfer of training effect for the stimuli sharing the color than those sharing the orientation with the trained stimulus. However, given the lack of significant results, we continued to examine this issue in the following experiments and review the overall results in the General Discussion.

There were two interesting findings in the eye movement results. First, we found that the proportions of the zero-saccade trials and the single on-target-saccade trials significantly increased after training, suggesting that the participants had a higher chance to identify the target before making any saccade. Second, we observed an increase in search initiation time after training for other correctly responded trials. This increase might reflect a change in search strategy as it could increase the probability of correct identification of target with fewer saccades. The potential mechanism that could account for these two findings was that training improved the covert attention to the trained stimulus. We will examine this interpretation in Experiment 5 where EEG signals were recorded.

## Experiment 2

In Experiment 2, we used a search array with a larger number of items (i.e., set size) that were smaller in size, to examine whether the observed training effects from Experiment 1 could be generalized to a different set of stimulus parameters. The overall area of the search array remained unchanged to keep the search window comparable across experiments.

## Method

### Participants

Fifteen right-handed naïve participants (12 females, age range = 19–28 years, mean age = 23.7 years) with normal or corrected-to-normal vision were recruited for the experiment.

### Apparatus

The apparatus in Experiment 2 was identical to Experiment 1.

### Stimuli

The stimuli in Experiment 2 were identical to Experiment 1, except that each search array consisted of 144 items (12 ×12) which were smaller in size (0.68° × 0.27°) (Figure 1B).

### Procedure

The general procedure in Experiment 2 was identical to Experiment 1. As the task would become more difficult with the increased set size and smaller search items, participants completed two practice blocks with 100 trials in each block to familiarize with the task before the pretest. In the pretest session, we tested three targets (r_90, r_180, and g_90) with 160 trials per target to get enough correct trials for the analysis. The target was always r_90 in eight training blocks with 300 trials in each block, keeping similar training time with Experiment 1.

### Data analysis

The data analysis was identical to Experiment 1. Trials with incorrect responses (24.97%), RTs that were faster than 200 ms (0.12%), and latency of first saccade smaller than 80 ms (7.85%), were excluded from analyses.

### Results

#### d’ and RTs

The behavioral results of Experiment 2 are shown in Figure 3A.

**Figure 3.**
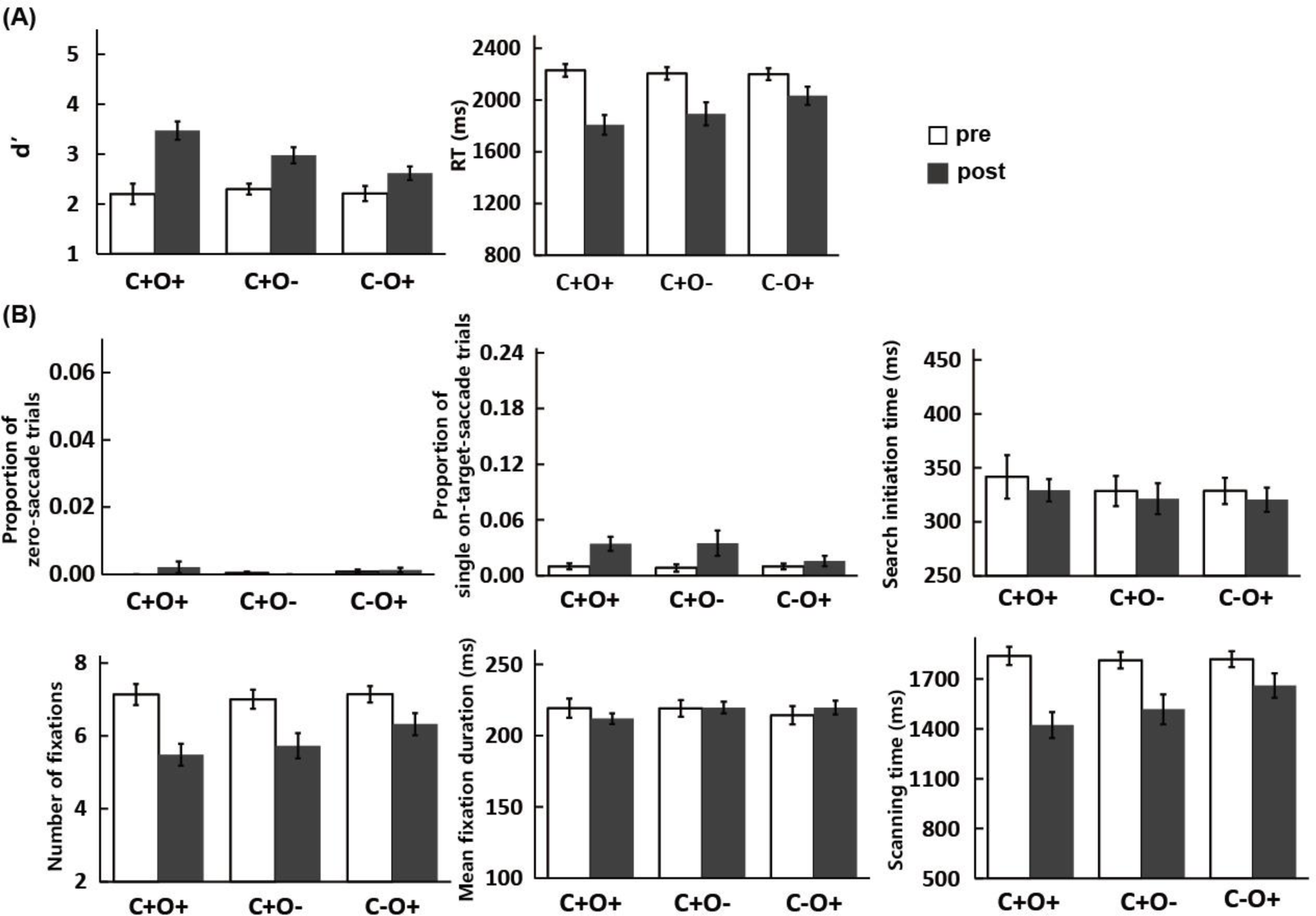
Results of Experiment 2. (A) Behavioral results for d’ and RTs. (B) Eye movement results for the proportion of zero-saccade trials, proportion of single on-target-saccade trials, and other trials’ search initiation time, number of fixations, mean fixation duration, and scanning time. Error bars represent standard errors of the mean across participants.

For d’, we observed significant effects on session (*F*(1,14) = 42.73, *p* < 0.001, *η^2^* = 0.75), target type (*F*(2,28) = 6.72, *p* < 0.01, *η^2^* = 0.32), and their interaction (*F*(1.29,18.12) = 9.40, *p* < 0.01, *η^2^* = 0.40). Simple-effects analysis (Bonferroni-corrected) revealed significant effects of training in C+O+ (*t*(14) = -5.91, *p* < 0.001, Cohen’s *d* = 1.53), C+O– (*t*(14) = -4.18, *p* < 0.001, Cohen’s *d* = 1.08), and C–O+ (*t*(14) = -3.66, *p* < 0.01, Cohen’s *d* = 0.94) conditions.

For RTs, we also observed significant effects on session (*F*(1,14) = 50.42, *p* < 0.001, *η^2^* = 0.78), target type (*F*(1.21,16.90) = 4.35, *p* < 0.05, *η^2^* = 0.24), and their interaction (*F*(1.24,17.42) = 9.03, *p* < 0.01, *η^2^* = 0.39). Simple-effects analysis (Bonferroni-corrected) revealed significant effects of training in C+O+ (*t*(14) = 9.27, *p* < 0.001, Cohen’s *d* = 2.39), C+O– (*t*(14) = 4.57, *p* < 0.001, Cohen’s *d* = 1.18), and C–O+ (*t*(14) = 3.52, *p* < 0.01, Cohen’s *d* = 0.91) conditions.

Paired t-tests on the transfer indices for C+O– and C–O+ conditions (Table 1) revealed a significant difference in RTs (*t*(14) = 2.55, *p* < 0.05, Cohen’s *d* = 0.66, TI was larger for C+O–) but not in d’ (*t*(14) = 1.66, *p* = 0.12, Cohen’s *d* = 0.43).

#### Eye movement

The eye movement results of Experiment 2 are shown in Figure 3B.

In contrary to Experiment 1, training did not significantly increase the proportion of the zero- saccade trials in Experiment 2. We did not observe significant effect on session (*F*(1,14) = 0.85, *p* =0.37, *η ^2^* = 0.06), target type (*F*(1.27,17.82) = 0.83, *p* = 0.40, *η ^2^* = 0.06), or their interaction (*F*(1.36,19.07) = 1.73, *p* = 0.21, *η^2^* = 0.11).

However, there was still a significant increase in the proportion of the single on-target-saccade trials in Experiment 2. We observed a significant effect on session (*F*(1,14) = 8.60, *p* < 0.05, *η^2^* = 0.38).

No significant effect was observed on target type (*F*(2,28) = 1.82, *p* = 0.18, *η^2^* = 0.12) or their interaction (*F*(1.39,19.42) = 2.03, *p* = 0.17, *η^2^* = 0.13).

For other correctly responded trials, training did not significantly increase the search initiation time. We did not observe significant effect on session (*F*(1,14) = 0.41, *p* = 0.53, *η^2^* = 0.03), target type (*F*(2,28) = 1.35, *p* = 0.28, *η^2^* = 0.09), or their interaction (*F*(2,28) = 0.09, *p* = 0.92, *η^2^* = 0.01).

Consistent with Experiment 1, training significantly reduced the number of fixations and scanning time and kept the mean fixation duration and verification time unchanged. For the number of fixations, there were significant effects on session (*F*(1,14) = 90.77, *p* < 0.001, *η^2^* = 0.87), target type (*F*(1.28,17.92) = 6.89, *p* < 0.05, *η^2^* = 0.33), and their interaction (*F*(2,28) = 7.32, *p* < 0.01, *η^2^* = 0.34). Simple-effects analyses (Bonferroni-corrected) revealed significant effects of training in C+O+ (*t*(14) = 10.45, *p* < 0.001, Cohen’s *d* = 2.70), C+O– (*t*(14) = 7.02, *p* < 0.001, Cohen’s *d* = 1.81), and C–O+ (*t*(14) = 4.06, *p* < 0.01, Cohen’s *d* = 1.05) conditions. For scanning time, there were significant effects on session (*F*(1,14) = 53.31, *p* < 0.001, *η^2^* = 0.79), target type (*F*(1.21,16.97) = 5.49, *p* < 0.05, *η^2^* = 0.28), and their interaction (*F*(1.44,20.12) = 10.44, *p* < 0.01, *η^2^* = 0.43). Simple-effects analyses (Bonferroni-corrected) revealed significant effects of training in C+O+ (*t*(14) = 9.10, *p* < 0.001, Cohen’s *d* = 2.35), C+O– (*t*(14) = 5.09, *p* < 0.001, Cohen’s *d* = 1.31) and C–O+ (t(14) = 3.19, *p* < 0.01, Cohen’s d = 0.82) conditions. The mean fixation duration and verification time were not significantly modulated by training (*ps* > 0.12), except that there was a significant interaction in the mean fixation duration. However, simple-effects analysis (Bonferroni-corrected) revealed no significant effect of training in C+O+ (*t*(14) = 1.48, *p* = 0.16, Cohen’s *d* = 0.38), C+O– (*t*(14) = -0.18, *p* = 0.86, Cohen’s *d* = 0.05), or C–O+ (t(14) = -1.61, *p* = 0.13, Cohen’s *d* = 0.42) condition.

Paired t-tests revealed no significant difference in transfer index between C+O– and C–O+ conditions in the number of fixations of the other correctly responded trials (*t*(14) = 1.93, *p* = 0.07, Cohen’s *d* = 0.50). There was a significant difference in scanning time between the two conditions (*t*(14) = 2.38, *p* < 0.05, Cohen’s *d* = 0.61).

### Discussion

With the increased set size and reduced stimulus size, we observed similar patterns of results in Experiment 2 as those in Experiment 1, except that no significant training effects for the proportion of the zero-saccade trials and the search initiation time of other correctly responded trials were observed. Apparently, increasing the set size in the search display and reducing the stimulus size made it less likely that the participants could locate the target when fixated at the center of the display. It was possible that the smaller item-to-item distance in Experiment 2 introduced crowding effect (Levi, 2008; Whitney & Levi, 2011) that prevented the participants properly perceived the items at their periphery visual field. To address this issue, in Experiment 3, we remained the stimulus size unchanged as in Experiment 2 but changed the set size to be the same as in Experiment 1, making the items in the search array less crowded than those in Experiment 2.

## Experiment 3

In Experiment 3, we kept the size of the search items identical to Experiment 2 but had the set size (6 × 6) to be the same as in Experiment 1 (Figure 1C). Importantly, this manipulation resulted in a less crowded search array as compared with both Experiment 1 and Experiment 2 while the overall search window remained unchanged, thus allowing us to investigate whether crowding effect was the key factor that contribute the different training effects between Experiments 1 and 2.

### Method

#### Participants

Fifteen right-handed naïve participants (eight females, age range = 18–24 years, mean age = 21.1 years) with normal or corrected-to-normal vision were recruited for the experiment.

#### Apparatus

The apparatus in Experiment 3 was identical to Experiment 1.

#### Stimuli

The stimuli in Experiment 3 were identical to Experiment 2, except that each search array consisted of 36 items (6 × 6).

#### Procedure

The general procedure in Experiment 3 was identical to Experiment 2.

#### Data analysis

The data analysis was identical to Experiment 1. Trials with incorrect responses (6.00%), RTs that were faster than 200 ms (0.01%), and latency of first saccade smaller than 80 ms (3.72%), were excluded from analyses.

### Results

#### d’ and RTs

The behavioral results of Experiment 3 are shown in Figure 4A.

**Figure 4.**
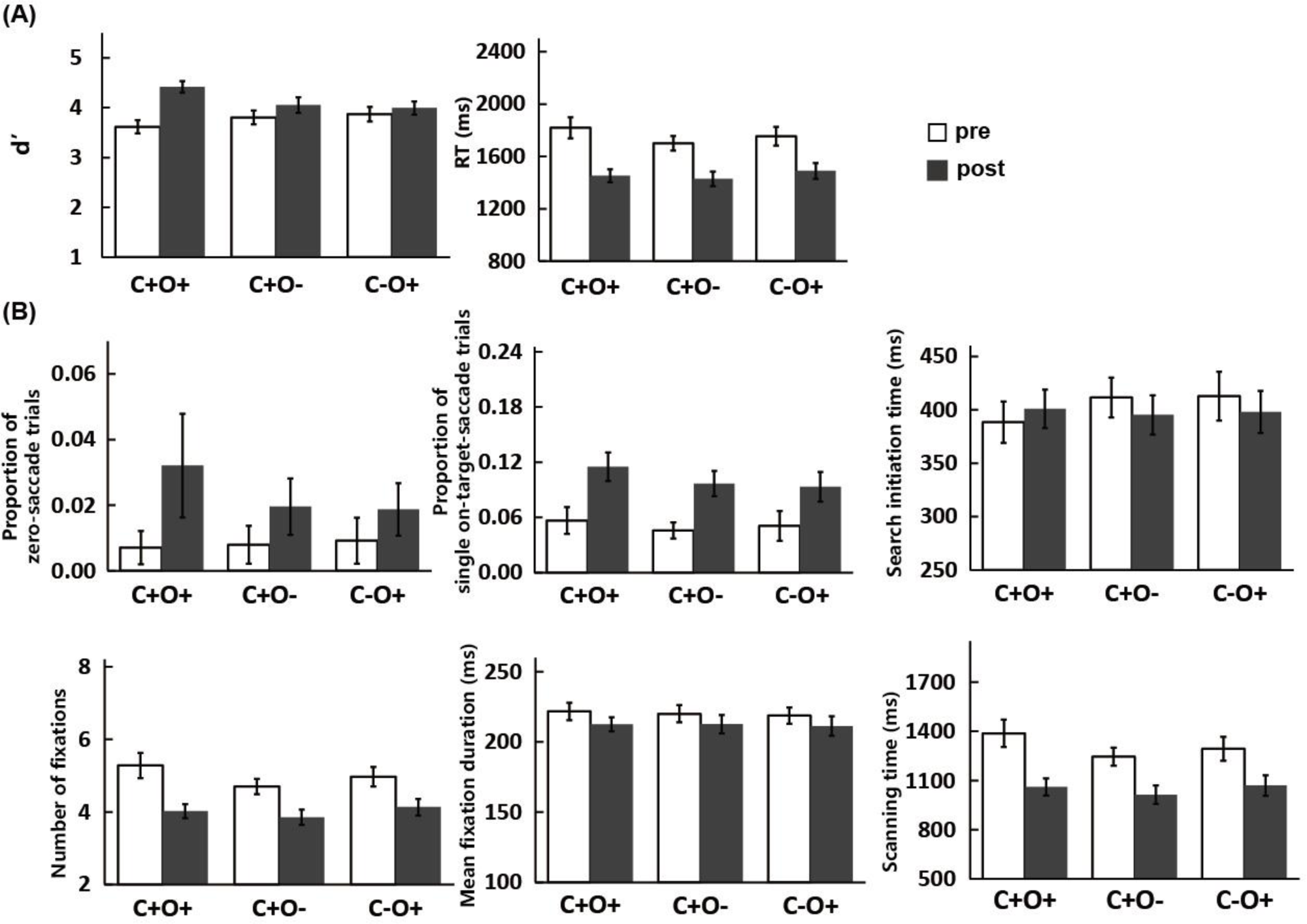
Results of Experiment 3. (A) Behavioral results for d’ and RTs. (B) Eye movement results for the proportion of zero-saccade trials, proportion of single on-target-saccade trials, and other trials’ search initiation time, number of fixations, mean fixation duration, and scanning time. Error bars represent standard errors of the mean across participants.

For d’, we observed significant effects on session (*F*(1,14) = 17.20, *p* < 0.001, *η^2^* = 0.55) and its interaction with target type (*F*(2,28) = 11.15, *p* < 0.001, *η^2^* = 0.44). No significant effect on target type was observed (*F*(2,28) = 0.38, *p* = 0.69, *η^2^* = 0.03). Simple-effects analysis (Bonferroni-corrected) revealed significant effect of training in C+O+ (*t*(14) = -7.27, *p* < 0.001, Cohen’s *d* = 1.88) condition. No significant effect of training was observed in C+O– (*t*(14) = -1.47, *p* = 0.16, Cohen’s *d* =0.38) or C–O+ (*t*(14) = -1.29, *p* = 0.22, Cohen’s *d* = 0.33) condition.

For RTs, we observed significant effects on session (*F*(1,14) = 62.40, *p* < 0.001, *η^2^* = 0.82) and target type (*F*(2,28) = 5.15, *p* < 0.05, *η^2^* = 0.27), but not their interaction (*F*(1.41,19.70) = 3.20, *p* = 0.08, *η^2^* = 0.19). Bonferroni-corrected post hoc tests showed that there were significant differences between C+O+ and C+O– conditions (*p* < 0.05). No significant difference was observed between C+O+ and C–O+ conditions (*p* = 1.00) and between C+O– and C–O+ conditions (*p* = 0.09).

Paired t-tests on the transfer indices for C+O– and C–O+ conditions found no significant difference in d’ (*t*(14) = 1.00, *p* = 0.33, Cohen’s *d* = 0.26) or RTs (*t*(14) = 0.61, *p* = 0.55, Cohen’s *d* = 0.16).

#### Eye movement

The eye movement results of Experiment 3 are shown in Figure 4B. The results were similar to Experiment 2.

For the proportion of the zero-saccade trials, there was no significant effect on session (*F*(1,14) = 3.90, *p* = 0.07, *η^2^* = 0.22), target type (*F*(1.37,19.19) = 0.61, *p* = 0.50, *η^2^* = 0.04), or their interaction (*F*(2,28) = 1.17, *p* = 0.32, *η^2^* = 0.08).

For the proportion of the single on-target-saccade trials, we observed a significant effect on session (*F*(1,14) = 40.16, *p* < 0.001, *η^2^* = 0.74). No significant effect was observed on target type (*F*(2,28) = 1.08, *p* = 0.35, *η^2^*= 0.07) or their interaction (*F*(2,28) = 0.29, *p* = 0.75, *η^2^* = 0.02).

For other correctly responded trials, training did not have significant effect on the search initiation time (session: *F*(1,14) = 0.46, *p* = 0.51, *η^2^* = 0.03; target type: *F*(2,28) = 0.81, *p* = 0.45, *η^2^* = 0.05; interaction: *F*(2,28) = 2.70, *p* = 0.08, *η^2^* = 0.16). For the number of fixations, there were significant effects on session (*F*(1,14) = 48.58, *p* < 0.001, *η^2^* = 0.78) and target type (*F*(2,28) = 10.62, *p* < 0.001, *η^2^*= 0.43), but not their interaction (*F*(2,28) = 3.09, *p* = 0.06, *η^2^*= 0.18). Bonferroni-corrected post hoc tests showed that there were significant differences between C+O+ and C+O– conditions (*p* < 0.01) and between C+O– and C–O+ conditions (*p* < 0.01). No significant difference was observed between C+O+ and C–O+ conditions (*p* = 0.81). For scanning time, there were significant effects on session (*F*(1,14) = 43.83, *p* < 0.001, *η^2^* = 0.76) and target type (*F*(2,28) = 8.22, *p* < 0.01, *η^2^*= 0.37), but not their interaction (*F*(1.42,19.92) = 2.79, *p* = 0.10, *η^2^* = 0.17). Bonferroni-corrected post hoc tests showed that there were significant differences between C+O+ and C+O– conditions (*p* < 0.01). No significant difference was observed between C+O+ and C–O+ conditions (*p* = 0.30) and between C+O– and C–O+ conditions (*p* = 0.10). For the mean fixation duration, there observed a significant effect on session (*F*(1,14) = 6.33, *p* < 0.05, *η^2^* = 0.31). There was no significant effect on target type (*F*(1.43,20.07) = 0.33, *p* = 0.65, *η^2^*= 0.02) or their interaction (*F*(2,28) = 0.13, *p* = 0.88, *η^2^*= 0.01).

Verification time remained unchanged after training (*ps* > 0.08).

Paired t-tests revealed no significant difference in transfer index between C+O– and C–O+ conditions in the number of fixations (*t*(14) = -0.99, *p* = 0.34, Cohen’s *d* = 0.26) or scanning time (*t*(14) = -0.81, *p* = 0.43, Cohen’s *d* = 0.21) for the other correctly responded trials.

#### Cross-experiment comparisons

To elucidate the influence of stimulus parameters on the training effects, we first examine the task difficulty across the three experiments by comparing their behavioral performance (d’ and RTs) at the pretest session. We performed mixed design ANOVAs on d’ and RTs of pretest session with target types (C+O+, C+O– and C–O+ conditions) as within-subject factor and experiment (Experiments 1, 2, and 3) as between-subject factor. For d’, the results revealed significant effects on experiment (*F*(2, 42) = 54.73, *p* < 0.001, *η^2^* = 0.72) and its interaction with target type (*F*(4, 84) = 2.99, *p* < 0.05, *η^2^* = 0.12). No significant effect was found on target types (*F*(2, 84) = 0.11, *p* = 0.89, *η ^2^* < 0.01). Simple-effects analysis (Bonferroni-corrected) revealed significant differences between Experiment 1 and Experiment 2 in all target types (C+O+: *t*(28) = 7.24, *p* < 0.001, Cohen’s *d* = 2.64; C+O–: *t*(28) = 9.18, *p* < 0.001, Cohen’s *d* = 3.35; C–O+: *t*(28) = 6.83, *p* < 0.001, Cohen’s *d* = 2.50) and between Experiment 2 and Experiment 3 in all target types (C+O+: *t*(28) = -5.76, *p* < 0.001, Cohen’s *d* = 2.10; C+O–: *t*(28) = -8.55, *p* < 0.001, Cohen’s *d* = 3.12; C–O+: *t*(28) = -7.84, *p* < 0.001, Cohen’s *d* = 2.86). No significant effect was observed between Experiment 1 and Experiment 3 in all types (*ps* > 0.34). For RTs, the results revealed significant effects on experiment(*F*(2, 42) = 31.88, *p* < 0.001, *η^2^*= 0.60) and target types (*F*(2, 84) = 3.58, *p* <0.05, *η^2^*= 0.08). No significant effect was found on interaction between the two factors (*F*(4, 84) = 1.12, *p* = 0.35, *η^2^* =0.05). Post hoc comparisons with Bonferroni correction revealed significant differences between Experiment 1 and Experiment 2 (*p* < 0.001) and between Experiment 2 and Experiment 3 (*p* < 0.001). We also observed significant difference between C+O+ and C+O– conditions (*p* < 0.05). No other significant effect was observed (*ps* > 0.15). These results suggest that the task difficulty was matched between Experiment 1 and Experiment 3, but the task difficulty of the two experiments was lower than that of Experiment 2.

Despite the difference in task difficulty, all three experiments showed significant behavioral training effects as measured with d’ and RTs. However, the three experiments showed different patterns of training effects in eye movements, particularly for the proportion of the zero-saccade trials (Exp. 1: *p* < 0.05, Exp. 2: *p* = 0.37, Exp. 3: *p* = 0.07) and the search initiation time of the other correctly responded trials (Exp. 1: *p* < 0.05, Exp. 2: *p* = 0.53, Exp. 3: *p* = 0.51). We performed three-way mixed design ANOVAs on these two measurements with session (pretest, posttest) and target types (C+O+, C+O– and C–O+ conditions) as within-subject factors and experiment (Experiments 1, 2, and 3) as between-subject factor. However, we did not observe meaningful between-experiment effect on both measurements, possibly due to the large number of factors and levels to be examined.

### Discussion

By equalizing set size between Experiment 1 and Experiment 3 and equalizing stimulus size between Experiment 2 and Experiment 3 (see Figure 1), we could examine the influence of these two stimulus parameters on the training effects.

First, the set size of the search array was likely to determine whether we could observe a significant training effect on the proportion of the zero-saccade trials. Within the same area of display, the set size of Experiment 2 was four times larger than those of Experiments 1 and 3, making the search items in Experiment 2 highly crowded. Meanwhile, we observed a significant training effect in Experiment 1 and a trend of significance in Experiment 3 for the proportion of the zero-saccade trials, while the effect in Experiment 2 was not significant. Because the proportion of the zero-saccade trials indicated how likely the participants could correctly identify the target without making saccades, this measurement would reflect the improvement of covert attention through training. The absence of increase in the proportion of the zero-saccade trials in Experiment 2 was likely the result of crowding effect as this effect has been shown to impair target identification at periphery (Kooi et al., 1994; Levi, 2008; Pelli & Tillman, 2008; Toet & Levi, 1992) and associate with the resolution of covert attention (Carrasco & Barbot, 2014; Montaser-Kouhsari & Rajimehr, 2005). Further, the higher task difficulty in Experiment 2 also agreed with this crowding interpretation. Taken together, the different patterns of training effect on the proportion of the zero-saccade trials could be interpreted as the results of different degrees of crowding in the search array between the experiments. That is, the participants in Experiment 2 were unlikely to improve their covert attention for the highly crowded search array.

Second, the stimulus size of the search array could also contribute to the training effect of the covert attention. The search initiation time of the other correctly responded trials demonstrated the contribution of covert attention in the correct identification that needed further verification with eye movements. Thus, it is a less deterministic measurement as compared with the proportion of the zero- saccade trials. By comparing Experiment 1 and Experiment 3, we found that the main difference between the two experiments was the search initiation time. These two experiments were matched in task difficulty and differed with each other only in the stimulus size. Therefore, we speculated that the smaller stimulus size made the participants in Experiment 3 harder to identify the target at periphery and showing no significant training effect on the search initiation time. This could be another supporting evidence for the role covert attention in the conjunction visual search training.

However, we would like to clarify that these proposals are still premature given the lack of direct statistical comparisons. The present results suggest that stimulus parameters such as set size and stimulus size had influenced the training effects that mostly related to covert attention in a free-viewing visual search paradigm. Future investigations with a specific design and a larger sample of participants are required to quantitatively examine the effects of various stimulus parameters.

## Experiment 4

In Experiment 4, we examined the practice effect that could be induced by the test sessions. We adopted the same procedure as in Experiment 2 except that there was no training session.

### Method

#### Participants

Fifteen right-handed naïve participants (11 females, age range = 18–24 years, mean age = 19.6 years) with normal or corrected-to-normal vision were recruited for the experiment.

#### Apparatus

The apparatus in Experiment 4 was identical to Experiment 1.

#### Stimuli

The stimuli in Experiment 4 were identical to Experiment 2 (Figure 1B).

#### Procedure

The procedure in Experiment 4 were identical to Experiment 2, except that there was no training in Experiment 4 (Figure 1E). After the pretest session, the participants had a two-hour break before the posttest session.

#### Data analysis

The data analysis was identical to Experiment 1. Trials with incorrect responses (31.78%), RTs that were faster than 200 ms (0.17%), and latency of first saccade smaller than 80 ms (6.68%), were excluded from analyses.

### Results

#### d’ and RTs

The behavioral results of Experiment 4 are shown in Figure 5A.

**Figure 5.**
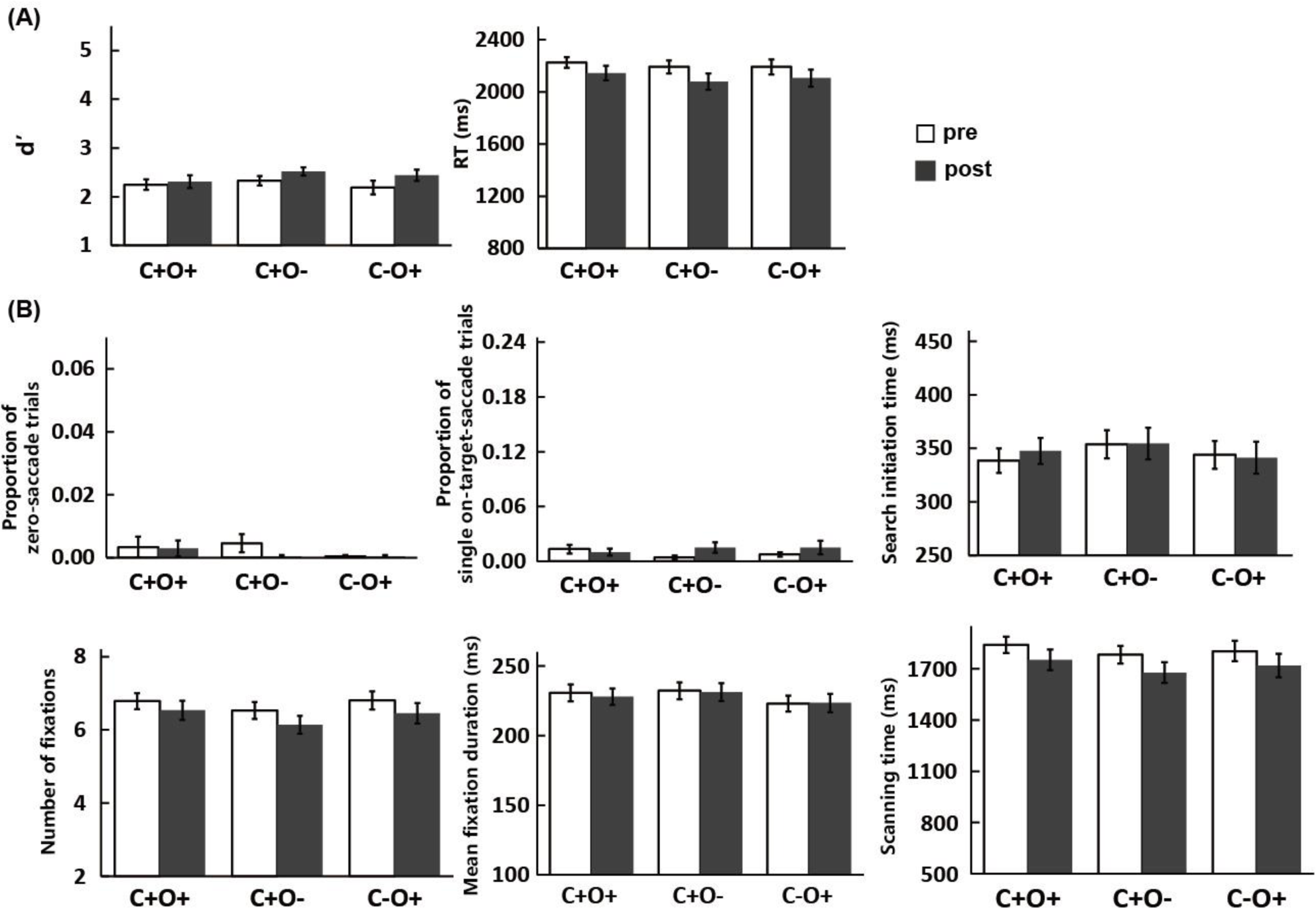
Results of Experiment 4. (A) Behavioral results for d’ and RTs. (B) Eye movement results for the proportion of zero-saccade trials, proportion of single on-target-saccade trials, and other trials’ search initiation time, number of fixations, mean fixation duration, and scanning time. Error bars represent standard errors of the mean across participants.

For d’, we observed significant effect on session (*F*(1,14) = 22.07, *p* < 0.001, *η^2^* = 0.61), but not on target type (*F*(2,28) = 1.32, *p* = 0.28, *η^2^* = 0.09) or their interaction (*F*(2,28) = 1.23, *p* = 0.31, *η ^2^* = 0.08).

For RTs, we observed significant effect on session (*F*(1,14) = 16.55, *p* < 0.01, *η^2^* = 0.54), but not on target type (*F*(2,28) = 1.23, *p* = 0.31, *η^2^* = 0.08) or their interaction (*F*(2,28) = 0.21, *p* = 0.81, *η ^2^* = 0.01).

We used a mixed design ANOVA to compare the improved d’ and RTs between Experiment 2 and Experiment 4. We calculated the improvement of d’ (posttest–pretest) and RTs (pretest–posttest) in two experiments separately. The mixed design ANOVA with target types (C+O+, C+O– and C–O+ conditions) and experiment (Experiment 2 and Experiment 4) as factors were conducted on improved d’ and RTs. For d’ improvement, there were significant effects on experiment (*F*(1, 28) = 24.15, *p* < 0.001, *η^2^*= 0.46), target type (*F*(1.52, 42.58) = 4.18, *p* < 0.05, *η^2^*= 0.13), and their interaction (*F*(1.52,42.58) = 10.21, *p* < 0.001, *η ^2^* = 0.27). Simple-effects analysis (Bonferroni-corrected) revealed significant effects of experiment in C+O+ (*t*(18.72) = 5.18, *p* < 0.001, Cohen’s *d* = 1.89) and C+O– (*t*(20.18) = 2.70, *p* < 0.05, Cohen’s *d* = 0.99) conditions. No significant effect of experiment was found in C–O+ condition (*t*(23.95) = 1.15, *p* = 0.26, Cohen’s *d* = 0.42). For RTs improvement, there were significant effects on experiment (*F*(1, 28) = 18.43, *p* < 0.001, *η^2^* = 0.40), target type (*F*(2, 56) = 4.99, *p* < 0.05, *η^2^*= 0.15), and their interaction (*F*(2, 56) = 5.12, *p* < 0.01, *η^2^*= 0.15). Simple-effects analysis (Bonferroni-corrected) revealed significant effect of experiment in C+O+ (*t*(28) = 5.48, *p* < 0.001, Cohen’s *d* = 2.00) and C+O– (*t*(28) = 2.51, *p* < 0.05, Cohen’s *d* = 0.92) conditions. No significant effect of experiment was found in C–O+ condition (*t*(28) = 1.38, *p* = 0.18, Cohen’s *d* = 0.50).

#### Eye movement

The eye movement results of Experiment 4 are shown in Figure 5B.

For the proportion of the zero-saccade trials, there was no significant effect on session (*F*(1,14)= 0.58, *p* = 0.46, *η^2^*= 0.04), target type (*F*(1.08,15.17) = 1.74, *p* = 0.21, *η^2^*= 0.11), or their interaction(*F*(2,28) = 0.66, *p* = 0.52, *η^2^* = 0.05).

For the proportion of the single on-target-saccade trials, there was no significant effect on session (*F*(1,14) = 1.40, *p* = 0.26, *η^2^* = 0.09), target type (*F*(2,28) = 0.22, *p* = 0.80, *η^2^* = 0.02), or their interaction (*F*(2,28) = 1.57, *p* = 0.23, *η^2^* = 0.10).

For the other correctly responded trials, training did not have significant effect on the search initiation time (session: *F*(1,14) = 0.18, *p* = 0.68, *η^2^* = 0.01; target type: *F*(2,28) = 3.07, *p* = 0.06, *η^2^* =0.18; interaction: *F*(2,28) = 0.67, *p* = 0.52, *η^2^* = 0.05). For the number of fixations, there were significant effects on session (*F*(1,14) = 14.70, *p* < 0.01, *η^2^* = 0.51) and target type (*F*(2,28) = 5.22, *p*< 0.05, *η^2^*= 0.27), but not their interaction (*F*(2,28) = 0.21, *p* = 0.81, *η^2^*= 0.01). Bonferroni-corrected post hoc tests showed that there were significant differences between C+O+ and C+O– conditions (*p* < 0.05). No significant difference was observed between C+O+ and C–O+ conditions (*p* = 1.00) and between C+O– and C–O+ conditions (*p* = 0.07). For scanning time, there was a significant effect on session (*F*(1,14) = 16.05, *p* < 0.01, *η^2^* = 0.53), but not on target type (*F*(2,28) = 2.19, *p* = 0.13, *η^2^* =0.14) or their interaction (*F*(2,28) = 0.07, *p* = 0.94, *η^2^* < 0.01). For the mean fixation duration, there was a significant effect on target type (*F*(2,28) = 3.85, *p* < 0.05, *η^2^* = 0.22). No significant effect onsession or their interaction was observed (*ps* > 0.63). For verification time, there was a significant interaction (*F*(2,28) = 3.43, *p* < 0.05, *η^2^* = 0.20). No significant effect on session or target type was observed (*ps* > 0.53). Simple-effects analysis (Bonferroni-corrected) revealed significant difference between pretest and posttest in C+O+ (*t*(14) = 2.33, *p* < 0.05, Cohen’s *d* = 0.60) condition. No significant effect was found in C+O– (*t*(14) = -1.04, *p* = 0.32, Cohen’s *d* = 0.27) or C–O+ (*t*(14) = 0.34, *p* = 0.74, Cohen’s *d* = 0.09) condition.

We used the mixed design ANOVA to compare the changes in the number of fixations and scanning time of the other correctly responded trials between Experiment 2 and Experiment 4. For number of fixations, we observed significant effects on experiment (*F*(1, 28) = 34.14, *p* < 0.001, *η^2^* =0.55) and its interaction with trial type (*F*(2, 56) = 4.60, *p* < 0.05, *η^2^* = 0.14). Simple-effects analysis(Bonferroni-corrected) revealed significant effects of experiment in C+O+ (*t*(28) = -5.61, *p* < 0.001, Cohen’s *d* = 2.05) and C+O– (*t*(28) = -3.94, *p* < 0.001, Cohen’s *d* = 1.44) conditions. There was a trend of significance in C–O+ condition (*t*(28) = -1.96, *p* = 0.06, Cohen’s *d* = 0.72). For scanning time, there were significant effects on experiment (*F*(1, 28) = 18.27, *p* < 0.001, *η^2^* = 0.39), target type (*F*(2, 56) =5.10, *p* < 0.01, *η^2^*= 0.15), and their interaction (*F*(2, 56) = 4.76, *p* < 0.05, *η^2^*

= 0.15). Simple-effects analysis (Bonferroni-corrected) revealed significant effects of experiment in C+O+ (*t*(28) = -5.06, *p* < 0.001, Cohen’s *d* = 1.85) and C+O– (*t*(28) = -2.67, *p* < 0.05, Cohen’s *d* = 0.98) conditions. No significant effect was observed in C–O+ condition (*t*(28) = -1.17, *p* = 0.25, Cohen’s *d* = 0.43).

### Discussion

Between-experiment comparisons in behavioral and eye movement results showed that training effects (Experiment 2) were significantly larger than practice effects without training (Experiment 4).

It should be noted that the training effects for the proportion of the zero-saccade trials and the search initiation time of other correct trials were absent in Experiment 2, making us unable to examine these two measurements in Experiment 4. However, for those significant training effects in Experiment 2, we demonstrated that they were not solely caused by practice. Therefore, we could conclude that our training protocols were effective despite that the whole experiment was completed within a single day.

## Experiment 5

In Experiment 5, we aimed to investigate the physiological signals that may contribute to the observed behavioral and eye movement training effects in Experiments 1, 2, and 3. We trained the participants with the same stimuli as in Experiment 1 and tested them with EEG and eye movement recorded after the training. Specifically, we examined the difference between conditions in which trained and untrained stimuli served as search target during the test. N2pc was initially identified as a key component that reflects the focusing of spatial attention onto the target location (Eimer, 2014; Luck & Hillyard, 1994). Following studies had further demonstrated that N2pc could also serve as a reliable indicator of attentional capture by physically salient stimulus (Hickey et al., 2006). In Experiment 5, we measured N2pc for the stimuli of interest (i.e., the trained target stimulus and an untrained stimulus) when they served as target or distractor during the conjunction visual search task. Specifically, we examined the traditional stimulus-locked N2pc, as well as the presaccadic (i.e., saccade-locked) N2pc that was suggested to indicate the covert attention before overt eye movement (Huber-Huber et al., 2016; Talcott & Gaspelin, 2021; Weaver et al., 2017).

### Method

#### Participants

Twenty right-handed naïve participants (nine females, age range = 18–26 years, mean age = 21.9 years) with normal or corrected-to-normal vision were recruited for the experiment.

#### Apparatus

All stimuli were displayed on a CRT monitor with a refresh rate of 60 Hz and spatial resolution of 1024 × 768 pixels. We controlled the stimulus presentation with Psychtoolbox 3.0 (Brainard, 1997; Pelli, 1997) in MATLAB programming environment (MathWorks, Natick, MA, USA). Participants were positioned 60 cm from the monitor in a dimly lit room. We used a chin rest to fix participants’ head position. EEG and eye movement data were simultaneously recorded. Eye movements were recorded using an EyeLink 1000 plus (SR Research, Ontario, Canada) eye tracker with a sampling rate of 1000 Hz. Gaze position was established using nine-point calibration and validation scheme. A drift correction was carried out before each block. EEG was digitized on-line at a sampling rate of 1000 Hz and recorded from using BrainAmp amplifier (Brain Products GmbH, Gilching, Germany) from a 64-channel cap (Easycap GmbH, Germany) with 64 Ag/AgCl electrodes arranged according to the International 10-20 System. The vertical electro-oculogram (EOG) was recorded below the right eye. Electrode impedances were kept below 10 kΩ during experiment.

#### Stimuli

The stimuli in Experiment 5 were identical to Experiment 1, except that g_45 could also serve as target during the training and test sessions.

#### Procedure

As shown in Figure 6A, each trial began with the presentation of a central fixation cross (presented for 450/500/550/600/650 ms) followed by a search array that appeared for 3000 ms or until key response. The inter-trial interval was 400/500/600 ms in practice and training sessions and 1400/1500/1600 ms in EEG test session to prevent the interference on EEG signal from the previous trial. These variations were introduced as a means of inter-trial jittering and to reduce the influence of participants’ expectation. We chose longer inter-trial interval in EEG session to reduce the potential interference of EEG signals between adjacent trials.

**Figure 6.**
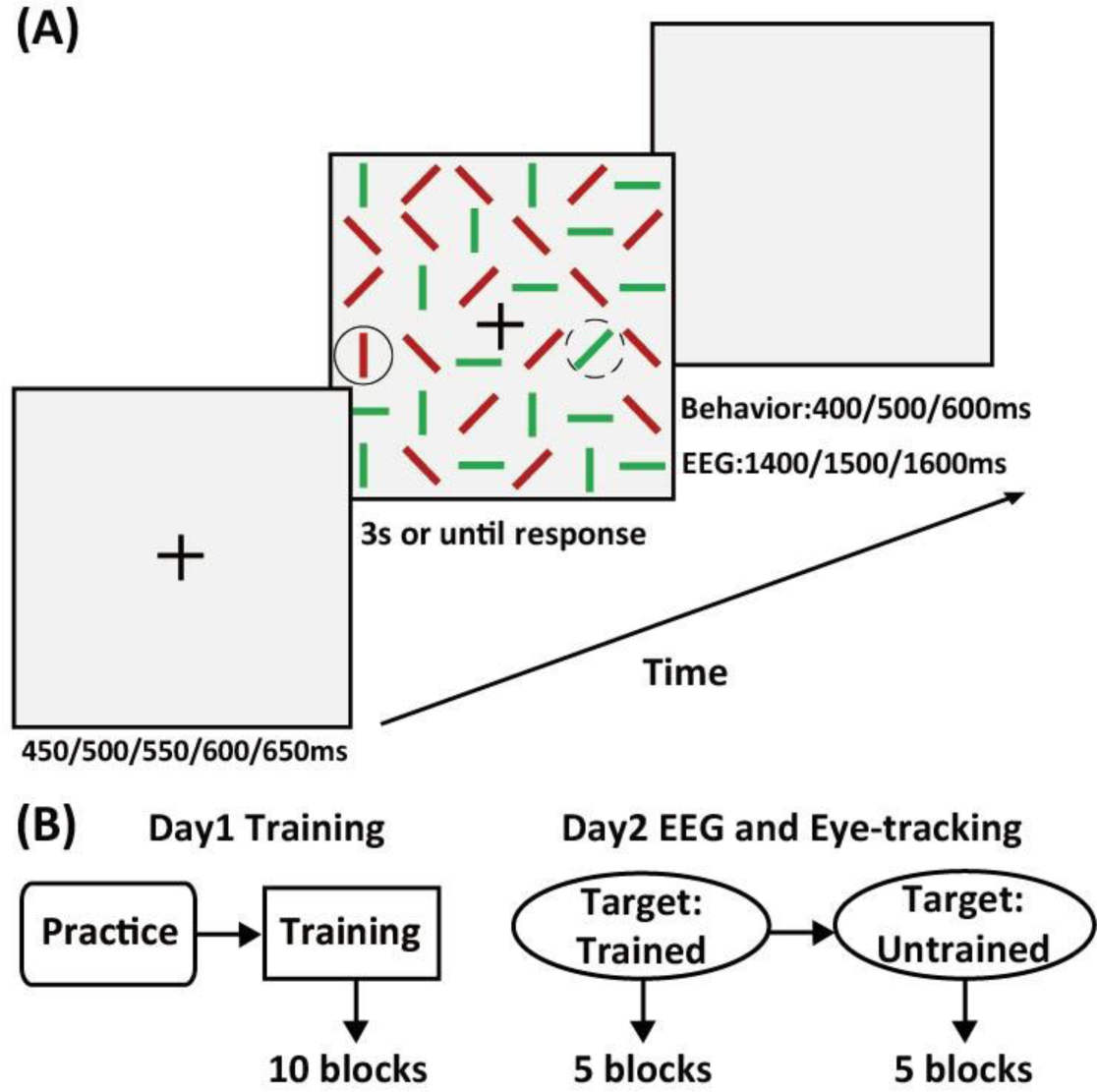
Experimental design of Experiment 5. (A) Stimulus and trial sequence. Solid line circle (trained target) and dashed line circles (untrained target served as distractor) are drawn for illustration purpose of training session and were not shown during the experiment. (B) Experimental procedure.

Participants completed the experiment in two days (Figure 6B). On the first day, they completed one practice block (120 trials with black and white line segments, target was black 90° line segments) to be familiar with the task and key response before training. During training (10 blocks with 300 trials in each block), participants were provided a break every 20 trials to reduce fatigue and boredom. On the second day, they completed 10 test blocks (120 trials in each block) with EEG and eye movements recorded.

During the training session, r_90 served as trained target for a half of participants and g_45 served as trained target for other half of participants. Also, g_0, g_90, r_45, and r_135 always served as distractors with an additional distractor being the untrained target (the target in other group of participants). In each search display, a half of items were in green color (g_0 and g_90 with equal numbers) and the other half of items were in red (r_45 and r_135 with equal numbers). The untrained target randomly replaced one of the distractors in the same color.

During test session with EEG recording, the trained target was tested in first five blocks where the untrained target was always presented as distractor, and the untrained target was tested in later five blocks where the trained target was always presented as distractor. Like the training session, there were equal number of green and red items in each search display. The untrained or trained target randomly replaced one of the original distractors in its color when it served as distractor in the block.

#### Data analysis

Analyses of behavioral and eye movement data were identical to Experiment 1 with two exceptions. First, paired t-tests between the trained target condition and untrained target condition during the test session were used to examine the training effects. Note that whether r_90 or g_45 served as the trained target or untrained target was counterbalanced across participants, we expected that difference between the trained target and untrained target conditions in the test session was due to the training. Second, we calculated the functional visual field for the zero-saccade trials and single on-target-saccade trials. Same trial exclusion criterion was applied in ERP analysis.

#### Functional visual field

We combined the zero-saccade trials (present trials only) and single on-target-saccade trials as a whole to calculate the FVF. The mean distance between the position of the initial fixation and the position of the target was used as the value of FVF. The calculated FVF could be considered to correspond the resolution and attentional FVFs in the literature that reflect the spatial scope of effective attentional processing (Wolfe, 2021; Wu & Wolfe, 2022). The difference of FVF between the trained and untrained conditions was considered the training effect. Due to a large number of participants that had no zero-saccade trials (present trials only) or single on-target-saccade trials in the pretest or posttest sessions, we did not perform the FVF analysis for Experiments 1 to 4.

#### EEG preprocessing

EEGLAB toolbox (v14.1.2; Delorme & Makeig, 2004) and EYE-EEG extension to EEGLAB (v0.81; Dimigen et al., 2011) in MATLAB environment were used for data analysis. All EEG data were re-referenced offline to the average of the left and right mastoids (TP9 and TP10) and then filtered with a 1 Hz high pass filter and a 40 Hz low pass filter. Eye tracking data were imported and synchronized with EEG signals. After synchronization, EEGs were segmented into epochs beginning at 100 ms before stimulus onset and ending at 600 ms after stimulus onset. Epochs that contained deteriorated eye tracking data (eye blinks or lost tracking) were removed (0.91% of all trials). Epochs that contained extreme EEG amplitudes (exceeded ±50 μV) of PO7 and PO8 channels were also removed (0.12% of all trials). Independent components were extracted from the combined EEG and eye movement data. The artifactual components were rejected based on the covariance with eye movement data using saccade to fixation variance ratio criterion of 1.1 (Plöchl et al., 2012). Baseline for event related potentials (ERPs) was set to be the mean voltage of 100 ms before stimulus onset.

#### Stimulus-locked N2pc

N2pc elicited by a specific item was defined as the differential ERP between contralateral and ipsilateral posterior scalp positions with respect to the item’s location. N2pc amplitudes were calculated for the four conditions defined by the corresponding item’s training history (trained vs. untrained) and role in the task (target vs. distractor) using the data of PO7 and PO8 electrodes. The two electrodes were averaged together to form an overall contralateral minus ipsilateral difference wave for each condition. To determine the period in which the averaged stimulus-locked N2pc was significantly deviated from baseline, successive t-test was used with a moving window of 20 ms (20 time points) in steps of 1 ms (1 time point). The criterion of the onset time of N2pc was that at least 40 consecutive windows (40 ms) reached significance at *p* < 0.05 level (Qu et al., 2017).

#### Saccade-locked N2pc

We identified and analyzed the N2pc in ERPs time-locked to the onset of the first saccade (Weaver et al., 2017). Trials with initial saccade latencies that below 80 ms (anticipations) or above 600 ms (retardations) were discarded. For the N2pc component locked to the first saccade, a criterion of at least 20 consecutive windows (20 ms) reached significance at *p* < 0.05 level was used to determine its onset time.

### Results

#### d’ and RTs

The behavioral results of Experiment 5 are shown in Figure 7A. Significant training effects were revealed when compared the d’ (*t*(19) = 5.38, *p* < 0.001, Cohen’s *d* = 1.20) and RTs (*t*(19) = -9.17, *p* < 0.001, Cohen’s *d* = 2.05) between the trained target and untrained target conditions.

**Figure 7.**
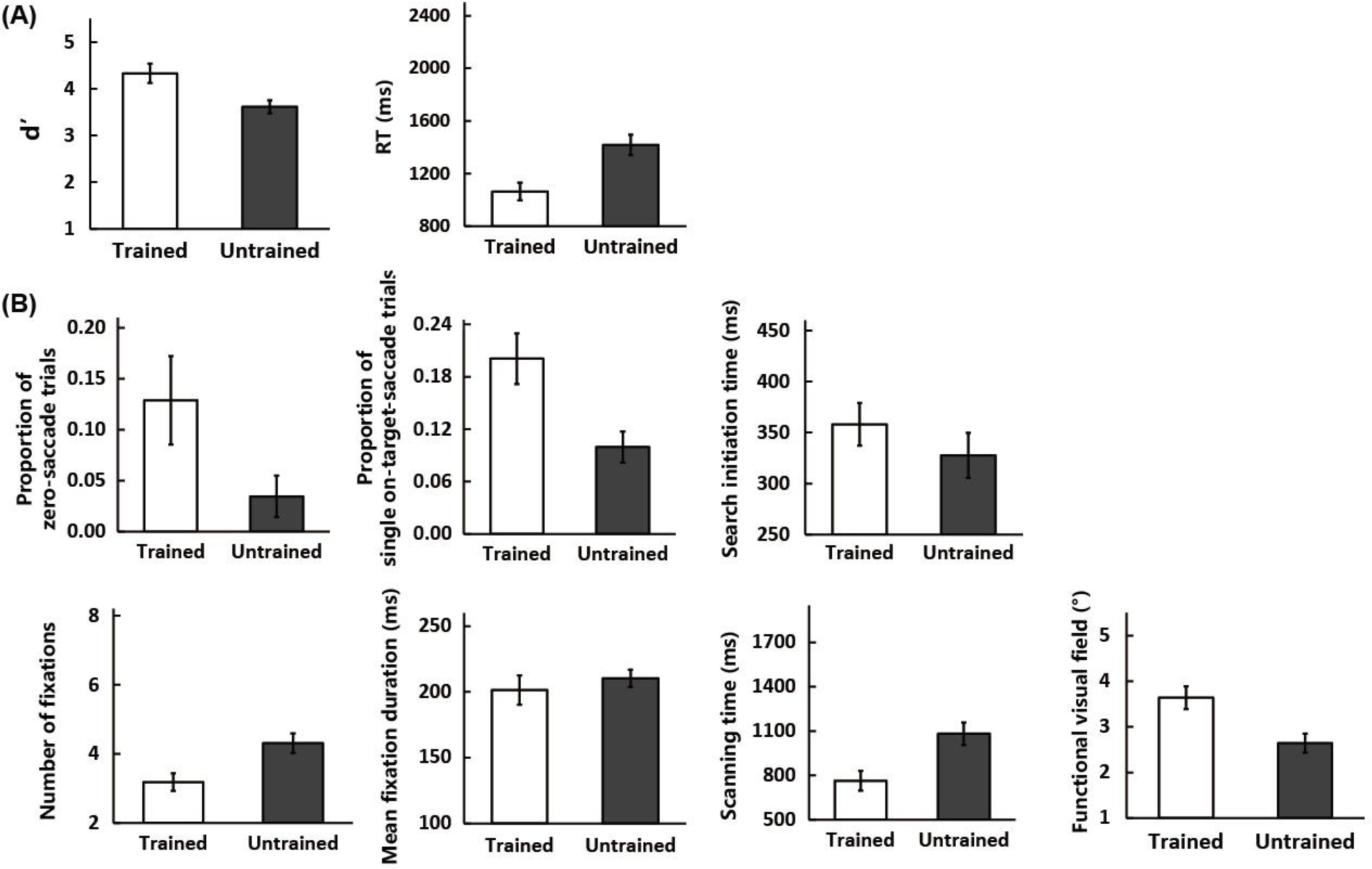
Behavioral and eye movement results of Experiment 5 for trials with trained and untrained targets in the post-training EEG session (day 2). (A) Behavioral results for d’ and RTs. (B) Eye movement results for the proportion of zero-saccade trials, proportion of single on-target-saccade trials, and other trials’ search initiation time, number of fixations, mean fixation duration, scanning time, and functional visual field. Error bars represent standard errors of the mean across participants.

#### Eye movement

The eye movement results of Experiment 5 also demonstrated similar effects as in Experiment 1 and are shown in Figure 7B.

The proportions of the zero-saccade trials (*t*(19) = 3.38, *p* < 0.01, Cohen’s *d* = 0.76) and single on-target-saccade trials (*t*(19) = 3.51, *p* < 0.01, Cohen’s *d* = 0.78) were significantly larger for the trained target condition as compared with the untrained target condition. For the other correctly responded trials, the training effects were significant for the search initiation time (*t*(19) = 2.42, *p* < 0.05, Cohen’s *d* = 0.54), number of fixations (*t*(19) = -9.47, *p* < 0.001, Cohen’s *d* = 2.12), and scanning time (*t*(19) = -9.31, *p* < 0.001, Cohen’s *d* = 2.08). No significant training effect was observed in mean fixation duration (*t*(19) = -1.36, *p* = 0.19, Cohen’s *d* = 0.30) or verification time (*t*(19) = -2.04, *p* = 0.06, Cohen’s *d* = 0.46) for these other trials.

Training also increased the functional visual field as indicated by the significantly larger distance between the initial fixation and the target location for the trained target as compared with the untrained target (*t*(19) = 3.83, *p* < 0.01, Cohen’s *d* = 0.86). The FVF was calculated based on the zero- saccade (present trials only) and single on-target-saccade trials. Therefore, we suggest that effective attentional field was enlarged after training and could contribute to the changes in the proportions of these two types of trials.

#### EEG

We examined the stimulus-locked and saccade-locked N2pc components for four conditions in which the trained and untrained stimuli served as target or distractor (TT: trained stimulus as target, TD: trained stimulus as distractor, UT: untrained stimulus as target, UD: untrained stimulus as distractor). The target-present trials were used in the analyses for N2pc components. Zero-saccade trials were analyzed as a separate condition for the stimulus-locked N2pc component.

For the stimulus-locked N2pc components (Figure 8A), we observed a significant N2pc when the trained stimulus served as target in the search task (TT: 234∼415 ms, peaked at 342 ms). No significant N2pc was found in other three conditions (i.e., TD, UT, and UD). Significant N2pc was also observed for the zero-saccade trials in the TT condition (288∼343 ms, peaked at 309 ms, 12 participants that had at least 10 zero-saccade trials in this condition were included in this analysis).

**Figure 8.**
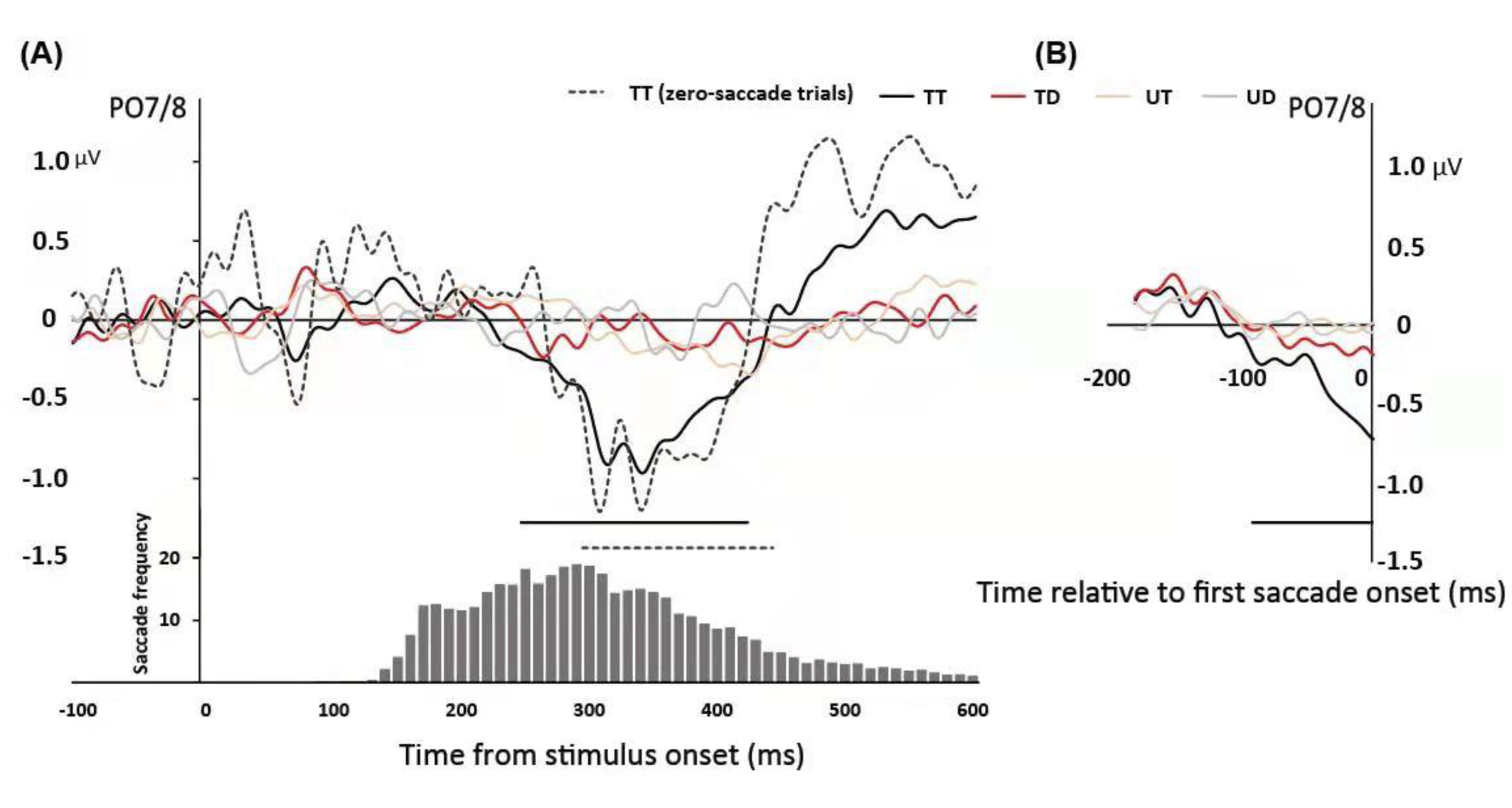
EEG results of Experiment 5. (A) Stimulus-locked N2pc components are shown for conditions in which the trained stimulus and untrained stimulus served as target or distractor. Black dash curve is the N2pc for TT condition of the zero-saccade trials. Frequency distribution of first saccade onset time aggregated across conditions and participants are presented below the ERP curves. (B) Saccade-locked N2pc components are shown for four conditions with eye movements. Periods of significant N2pc amplitudes were shown as straight lines below the ERPs. TT: trained stimulus served as target; TD: trained stimulus served as distractor; UT: untrained stimulus served as target; UD: untrained stimulus served as distractor.

For the saccade-locked N2pc components (Figure 8B), significant N2pc was only evident when the trained stimulus served as target (TT: -94∼0 ms). No significant saccade-locked N2pc was found in other three conditions.

### Discussion

The behavioral and eye movement results demonstrated significant training effects when comparing the two conditions in which the trained and untrained stimulus served as the search target, respectively. These results were comparable to the observed effects in Experiment 1. Particularly, the eye movement results revealed an enlarged functional visual field that was associated with the increased scope of effective attentional processing after training.

The EEG results revealed both stimulus-locked and saccade-locked N2pc components that were elicited by the trained stimulus when it served as target in the test session. Specifically, the untrained stimulus shared no feature with the trained target and did not elicit significant N2pc component. The significant stimulus-locked N2pc component for the trained target condition, especially for those of the zero-saccade trials, suggest an enhanced covert attention to the trained target. A recent study has shown that the presaccadic N2pc component could be observed only if covert attention was allocated to the stimulus before the generation of the eye movement (Talcott & Gaspelin, 2021). Hence, the saccade-locked N2pc could also be explained as the enhanced covert attention to the trained target. Taken together, we suggest that the increased covert attentional processing to the trained stimuli was the underlying mechanism that drove the behavioral and the overt eye movement changes after training.

### General Discussion

The present study adopted a conjunction visual search task and examined the effects of single-day training on the task performance. We measured eye movements and EEG signals to elucidate the potential mechanisms underlying the training effects. The results showed that training on the conjunction visual search task led to significant behavioral effects on d’ and RTs (Experiments 1, 2, 3, and 5) and such improvement was significantly reduced when training was not taking place (Experiment 4). Meanwhile, the eye movement data revealed that the behavioral improvement was accompanied with reduced fixation number in all training protocols (Experiments 1, 2, 3, and 5) and increased proportion of zero-saccade trials and search initiation time of other correctly responded trials only in Experiments 1 and 5. This between-experiment difference could be attributed to the stimulus’ parameters that varied between them. Finally, we identified stimulus-locked and saccade-locked N2pc components elicited by the trained target stimulus as the neural signatures for training-induced enhancement in covert attention (Experiment 5). These findings collectively suggest that perceptual training on visual search increases the attentional priority of the trained stimuli and facilitates behavioral performance through optimized eye movement patterns.

In our experiments, we adopted a free viewing paradigm for the conjunction visual search task and were able to examine the patterns of eye movements in addition to the traditional behavioral measurements (Baluch & Itti, 2010). Free viewing is a natural way of observation when our visual system functions in various visual tasks. Allowing participants to move their eyes during the search tasks was suggested as a key factor for understanding the mechanism of selective attention in complex natural environment (Hayhoe & Ballard, 2005). Importantly, we found significant behavioral improvement after training that was consistent with previous studies that required participants keeping fixation during the task (An et al., 2012; Bueichekú et al., 2016, 2019; Clark et al., 2015; Leonards et al., 2002; Qu et al., 2017; Reavis et al., 2016, 2018; Sigman & Gilbert, 2000; Sirenteanu & Rettenbach, 2000; Sireteanu & Rettenbach, 1995; Su et al., 2014). These results suggest that the free viewing paradigm is a valid approach to investigate visual search training.

Consistent with previous studies, we found improved d’ and reduced RTs, as well as the transfer of training effects to stimuli that shared a feature with the trained target. The transfer effect was evident in previous investigations that showed non-specific training effect of visual search task (Lee et al., 2018; Sirenteanu & Rettenbach, 2000; Sireteanu & Rettenbach, 1995; Su et al., 2014), suggesting that a feature-based attention enhancement mechanism rather than a unitization mechanism was adopted during the training. Specifically, we calculated transfer indices of the dependent variables for Experiments 1, 2, and 3. For these transfer indices, we were interested in the comparisons between the C+O– and C–O+ conditions. That is, whether the transfer of training effects was equal between the color (C+O–) and orientation (C–O+) features. We found significantly larger transfer indices for the C+O– as compared with C–O+ condition in RTs and scanning time of Experiment 2. There were also trends of significance in the same direction in RTs of Experiment 1 and number of fixations of Experiment 2. Despite other comparison were not significant, most of the dependent variables (13 out of 15) in Table 1 showed larger transfer indices for the C+O– condition. These results imply that the transfer of training effect was larger for the color feature than the orientation feature. This is consistent with previous literature that color has higher priority for processing than other features in the visual system when used in conjunction with other features (Luria & Strauss, 1975; Williams & Reingold, 2001; Williams, 1966) .

Our analyses were based on correct trials and these trials were assigned to three groups. Among them, the zero-saccade trials had no saccade during the search. In the other two groups, the single on-target-saccade trials and other correctly responded trials, at least one saccade was executed during the search period. In Experiment 5, we observed larger proportion of zero-saccade correct trials for the trained than untrained target. This difference was accompanied by a significant stimulus-locked N2pc for the trained target, indicating stronger covert attention in this condition. For the single on-target-saccade trials and other correctly responded trials, the search initiation time was longer for the trained than untrained target. The observed difference in these saccadic trials was accompanied by a significant saccade-locked N2pc for the trained target. This presaccadic N2pc component has been suggested to reflect the covert attention before the generation of the eye movement (Talcott & Gaspelin, 2021; but see Li et al., 2021 for potential dessociation of presaccadic and covert attention). Additionally, we also revealed an enlarged functional visual field after training for the zero-saccade trials and single on-target-saccade trials. The increased scope of effective attentional processing could also contribute to the improved detection of target without saccades. Therefore, we could conclude that training enhanced the covert attention to the trained target stimulus, at least in the zero-saccade trials. This enhancement led to either larger proportion of trials in which the participants could identify the target while fixated the central fixation or increased search initiation time. Specifically, though increasing search initiation time showed apparent detrimental effect on search speed, it nevertheless provided the opportunity to identify the target without further eye movements that could greatly save the search time. The results in Experiment 5 suggest that training enhanced covert attentional processing towards the training target stimulus and could underlie the changed overt eye movement patterns.

Previous studies found that the training effect could also be observed while the trained target served as distractors (Qu et al., 2017), an effect that was not observed in our experiment. There were two possibilities that may account for the difference between the studies. First, the tasks and stimuli were different in the two studies. We used a conjunction visual search task with color and orientation as the critical features. In Qu et al. (2017), shape served as the only critical feature in the visual search task. Second and more importantly, we adopted a short training protocol that was completed within a single day, whereas the participants in Qu et al. (2017) were trained for several days. More extensive training is likely to result in stronger attentional bias to the trained stimulus and this elevated attentional processing could facilitate its competition against the search target for attentional selection.

In summary, with a free-viewing conjunction visual search training paradigm, we found significant improvement in behavioral performance after training. The behavioral training effects were accompanied by reduced number of saccades, as well as increased proportion of zero-saccade trials and search initiation time that were influenced by stimulus parameters. EEG results showed both stimulus-locked and saccade-locked N2pc components when the search target was the trained one, indicating training-induced enhancement in covert attention. Taken together, our findings offered new insights that the combination of the enhanced covert attention to target and optimized overt eye movements contribute to the behavioral improvement after visual search training.

